# Transgenic human dipeptidyl peptidase-4 Syrian hamsters support MERS coronavirus infection and contact transmission

**DOI:** 10.64898/2026.05.26.725122

**Authors:** Tong Wang, Yanan Liu, Rong Li, Nathan Merrill, Zhongde Wang, Peter J. Halfmann

## Abstract

Middle East respiratory syndrome coronavirus (MERS-CoV) is a global health concern due to a high fatality rate associated with human infections and no approved vaccines or therapeutics. While Syrian hamsters are a value animal model for coronavirus research, including SARS-CoV-2, MERS-CoV does not infect wild-type hamsters. Here, we generated transgenic Syrian hamsters expressing human dipeptidyl peptidase-4 (hDPP4), the cellular receptor for MERS-CoV., MERS-CoV replicated efficiently in the respiratory tract tissues of hDPP4 hamsters, causing lethal disease. Treatment with the 3CLpro inhibitor nirmatrelvir significantly reduced viral titers in the lower respiratory tract of infected hDPP4 hamsters. While airborne transmission was not observed, direct contact transmission was observed in all contact hDPP4 hamsters cohoused with infected cage mates. Immunization with purified MERS receptor-binding domain protein reduced virus replication and disease severity but did not prevent direct contact transmission. Collectively, our findings demonstrate that hDPP4 transgenic Syrian hamsters are useful for studying MERS-CoV pathogenesis, transmission, and countermeasure efficacy.

## Introduction

Middle East respiratory syndrome coronavirus (MERS-CoV) emerged in 2012 and continues to pose a zoonotic and public health threat due to its high case fatality rate and sporadic outbreaks across the Middle East and beyond^1^. Moreover, no licensed vaccines or antiviral therapies are available for MERS-CoV infection. As of August 2025, the World Health Organization has reported 2627 laboratory-confirmed cases of MERS-CoV infection worldwide, including 947 deaths, which yields a case-fatality ratio (CFR) of 36% (WHO website, MERS situation update, August 2025). For SARS-CoV-2, experimental transmission models have been well established, primarily using Syrian hamsters and ferrets^2-4^. However, efficient MERS-CoV transmission has not been robustly demonstrated in a practical small animal model, limiting experimental study of transmission dynamics. Establishing such a model would advance both our mechanistic understanding of MERS-CoV transmission and the preclinical evaluation of candidate interventions.

Conventional laboratory animals such as mice and hamsters are not susceptible to MERS-CoV due to incompatibility between the viral spike (S) protein and their endogenous dipeptidyl peptidase-4 (DPP4, also known as CD26) receptor^5-8^. Several animal species have been evaluated for MERS-CoV susceptibility, including mice, ferrets, rabbits, and non-human primates, but each has limitations. Wild-type mice are refractory to infection^9^, while human DPP4 (hDPP4) transgenic and adenovirus-transduced mouse models often show limited respiratory tropism or develop neurologic symptoms that are not characteristic of human disease^10-12^. Ferrets and rabbits support only limited or transient viral replication in the upper airways without severe disease^13,14^. Nonhuman primates (e.g., rhesus macaques and common marmosets) develop respiratory illness and serve as valuable pathogenesis models, but their high cost and limited availability restrict their use in large studies^15-17^. To date, no small animal model has demonstrated efficient and reproducible transmission of MERS-CoV, a major obstacle to the study of viral spread and the development of effective countermeasures. There is an urgent need for a genetically tractable, physiologically relevant, and transmissible small animal model that supports robust respiratory replication and enables evaluation of both viral pathogenesis and antiviral therapeutics.

DPP4 is widely expressed in human tissues, including the respiratory epithelium, kidney, liver, and immune cells, and serves as the functional entry receptor for MERS-CoV^5^. To overcome species-specific receptor restriction, we generated a transgenic Syrian hamster line expressing human DPP4 (hDPP4), thereby enabling efficient viral entry and replication. Upon intranasal inoculation with MERS-CoV, hDPP4 hamsters developed high viral titers throughout the respiratory tract and sustained replication through six days after infection. Using this model, we characterized viral replication kinetics, assessed transmission dynamics, and evaluated the therapeutic efficacy of vaccination and antiviral treatment. Our findings establish the hDPP4 transgenic Syrian hamster as a relevant small animal model for investigating MERS-CoV pathogenesis, transmission, and medical countermeasure development.

## Results

### Transgenic expression of human dipeptidyl peptidase 4 (hDPP4) in Syrian hamsters

To generate transgenic hamsters expressing hDPP4, we constructed a PiggyBac vector, pmhyGENIE-3-K18-hDPP4 (Fig. 1a), in which the expression of the hDPP4 cDNA is under the control of the human cytokeratin 18 promoter and other regulatory sequences. Pronuclear (PN) injection of this vector into hamster zygotes resulted in the production of 12 F0 hamsters. Genomic PCR genotyping analysis with three overlapping primer pairs (Table 1), F11/R11, F26/R26, and F31/R31, covering the entire K18-hDPP4 transgenic cassette, identified four hDPP4-positive F0 founders: F0M1, F0M7, F0M9, and F0F12 (Fig. 1b). We chose founder F0F12 to establish a hDPP4 transgenic breeding colony by first breeding it with a wild type (WT) male hamster to generate F1 offspring and subsequently intercrossing the hDPP4-positive F1 littermates. Animals produced from this line show normal development and fertility.

**Figure 1.**
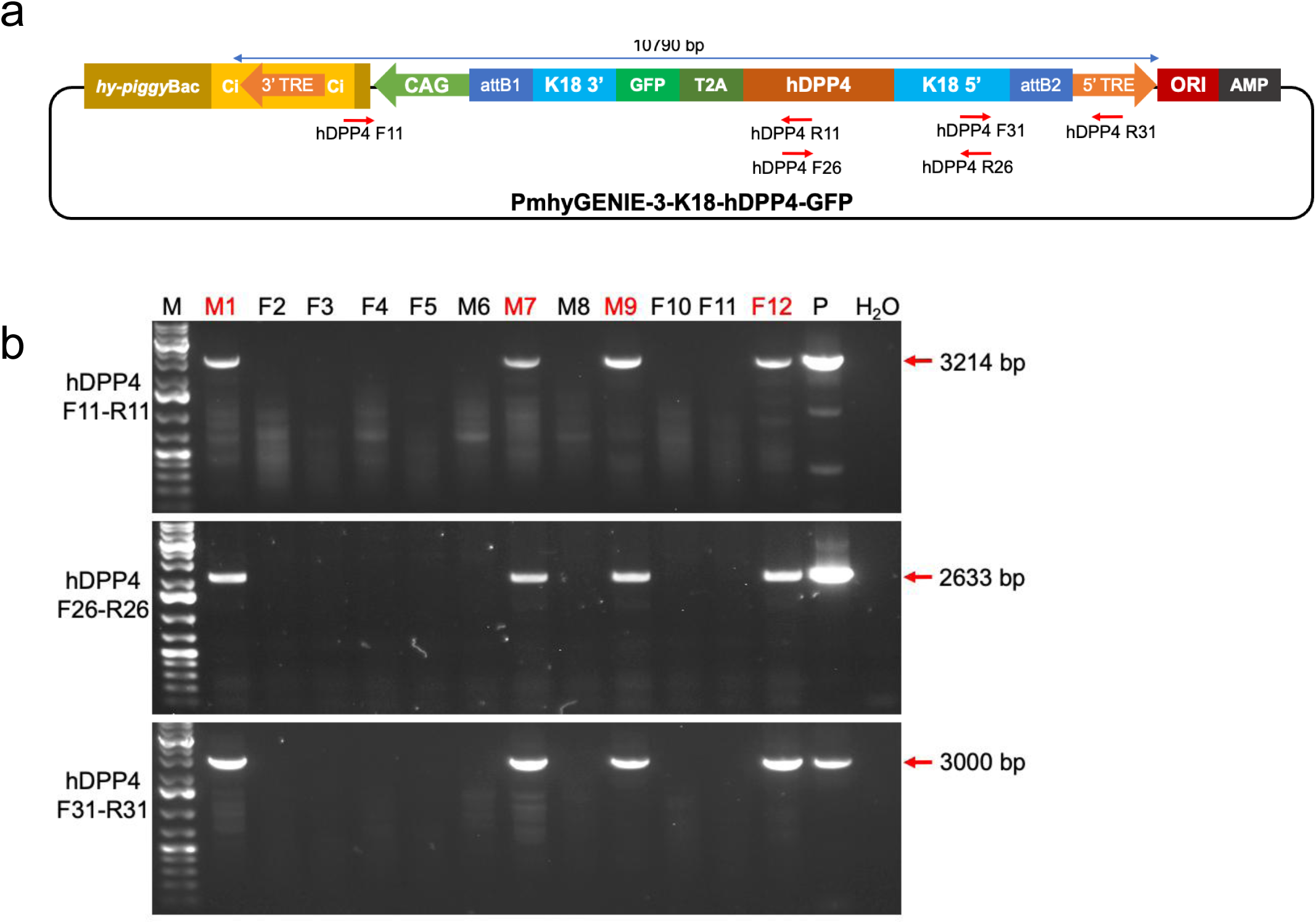
Generation of transgenic hDPP4 golden Syrian hamsters through piggyBac-mediated transgenesis. **a** Schematic representation of the piggyBac vector, pmhyGENIE-3-K18-hDPP4-GFP. The transposon cassette for genomic integration contains the K18 5’ region, human DPP4 CDS, T2A linker, GFP (as a reporter to visualize hDPP4 expression), K18 3’ region, and poly(A) signal, all flanked by the 3′- and 5′ terminal repeat element (TRE), and a hyperactive transposase (hy-piggyBac) driven by the CAG promoter. The chimeric intron (Ci) is located upstream of hy-piggyBac to enhance its expression. The positions of primers for genotyping transgenic pups are indicated as red arrows. The transposon is indicated as a blue line with a double-headed arrow. Diagrams are not to scale. **b** PCR genotyping the 12 F0 hamsters. F0M1, F0M7, F0M9, and F0F12 were identified as transgenic, and the rest of the pups were wild type. M, 1 kb Plus DNA Ladder; P, 5 ng of pmhyGENIE-3-K18-hDPP4-GFP plasmid used as positive control; H_2_O, H_2_O served as a negative control.

**Table 1:**
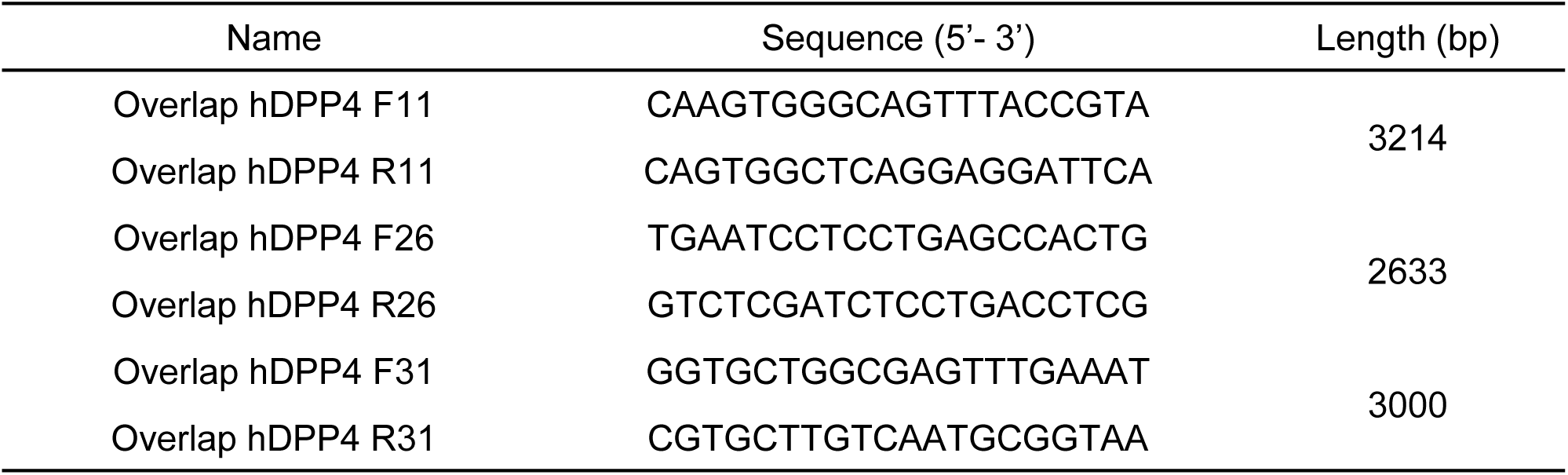
Primers used for genotyping transgenic hDPP4 hamsters.

To determine the genomic integration site(s) of the piggyBac transposon and the K18-hDPP4 transgene cassette, Cergentis (Utrecht, The Netherlands) performed Targeted Locus Amplification (TLA) analysis on genomic DNA isolated from F0F12. TLA sequencing identified five independent genomic integration sites in hamster genomic scaffolds (i.e., KB708127.1, KB708154.1, KB708173.1, KB708195.1 and KB708199.1) in this founder animal. We named these genomic integration sites TLA1, TLA2, TLA3, TLA4, and TLA5 (Table 2). No DNA sequence outside the terminal repeat elements (TREs) regions of the piggyBac transposon was detected, confirming each of the piggyBac transposon integration events was through the cut-and-paste mechanism mediated by TRE/transposase as designed. As the K18-hDPP4 transgene cassette at each of the genomic integration sites in F0F12 may pass on to next generations independently from breeding (based on the 5′ and 3′ junction sequences identified by TLA), junction-specific PCR primers covering the unique genomic junctions generated by the integration of the piggyBac transposon in the hamster genome were designed to detect the presence of the K18-hDPP4 transgene cassette at each of the integration sites in F1 and the subsequent filial generations of hamsters (Table 2). PCR primers were also designed to identify the WT alleles corresponding to each of the integration sites in the subsequent filial generations of hamsters derived from F0F12 (Table 2). Due to the size of the piggyBac transposon (>10 kb) and our PCR conditions, only WT alleles can be amplified with WT primers (Fig. 2a). Genotyping of the integration sites in a litter of F2 generation animals with junction-specific PCR for each of the TLA sites (TLA1–5) revealed that hamsters F2M4 and F2M9 carry the hDPP4 transgene at the KB708127.1 integration site only (Fig. 2b; junction PCRs for the other TLA sites are not shown). PCR analysis with WT primers further revealed that both F2M4 and F2M9 were homozygous for the hDPP4 transgene cassette (WT PCR negative), while all remaining hDPP4-positive animals were heterozygous (WT PCR positive) for this locus (Fig. 2b). We further crossed these two homozygous hamsters with their heterozygous littermates and conducted genotyping analysis with both junction PCR primers and WT PCR primers to identify homozygous hamsters. These homozygous hamsters were then used as breeders to establish a hDPP4 transgenic hamster colony. Therefore, from founder F0F12, we established a hDPP4 transgenic hamster line that carries one copy of the K18-hDPP4 transgenic cassette integrated in scaffold KB708127.1 of the hamster genome. All infection experiments described herein were performed using this hDPP4 transgenic line.

**Figure 2.**
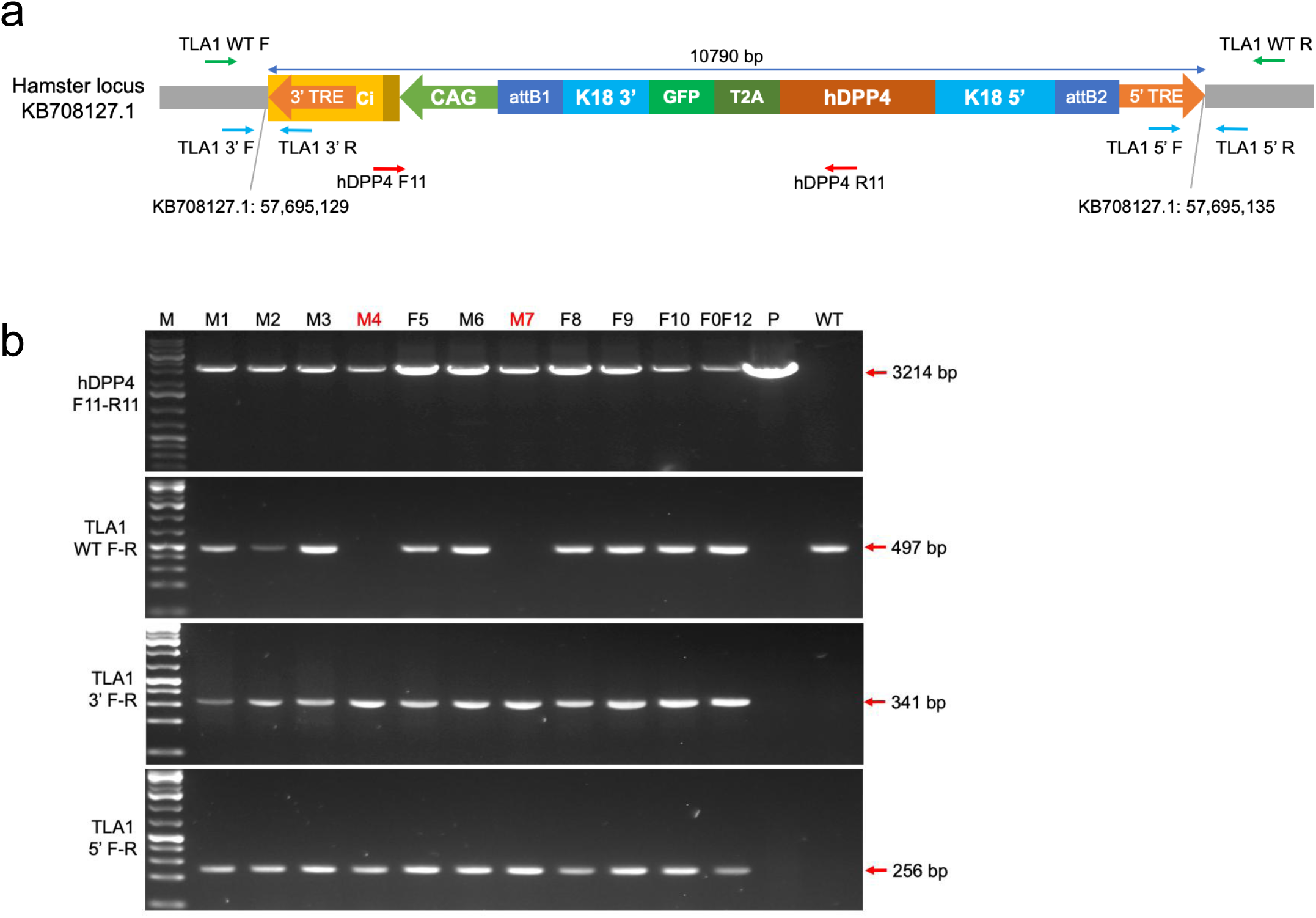
Genotyping of hDPP4 transgenic animals with TLA-specific PCRs. **a.** Schematic diagram of the hDPP4 transgene integration locus on scaffold KB708127.1 (TLA1). PCR primers used to validate the 5’ and 3’ genomic-transgene junction integration sites and wild type alleles are indicated as blue and green arrows, respectively. PCR primers used to detect part of the transgenic cassette are indicated as red arrows **b** PCR genotyping results of a litter of F2 pups produced from intercrossing of F1 hDPP4-positive hamsters. All 10 of the F2 pups were identified as hDPP4-positive with primer pair F11–R11 and junction primers TLA1 3’F-R and TLA1 5’F-R. Among these 10 pups, all except for F2M4 and F2M9, were PCR positive with the TLA1 WT F-R primers, indicating that F2M4 and F2M9 are homozygous for the hDPP4 transgene at TLA1, whereas the others are heterozygous. Junction PCRs for TLA2, TLA3, TLA4, and TLA5 in each of these animals were negative (data not shown), revealing that these TLAs were separated from TLA1 through breeding and that these animals only carry the hDPP4 transgene in the TLA1 locus. M, 1 kb Plus DNA Ladder; P, 5 ng of pmhyGENIE-3-K18-hDPP4-GFP plasmid; WT, gDNA from a wild type hamster; F0F12, gDNA from founder F0F12.

**Table 2:**
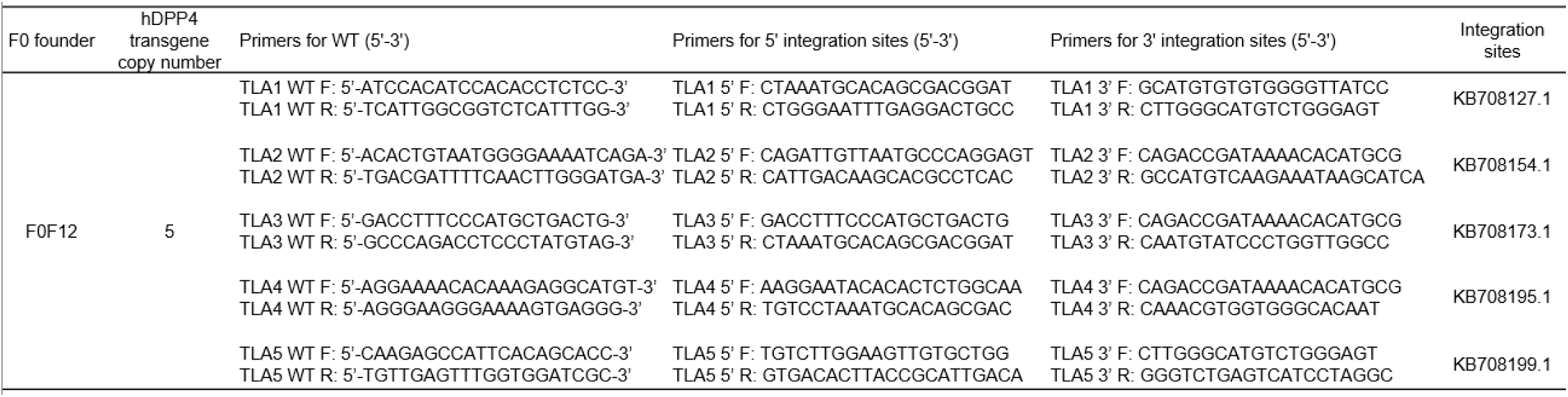
Primers for TLA analyses of pmhyGENIE-3-K18-hDPP4 integration.

### Efficient replication of MERS-CoV in the respiratory tissues of hDPP4 transgenic Syrian hamsters

To confirm the susceptibility of the hDPP4 hamsters to infection with MERS-CoV, we intranasally inoculated transgenic animals (n=4, males, 4–5 months old) with virus (MERS-CoV EMC/2012 strain; 30 µl total volume) at a dose of 10⁵ plaque-forming units (pfu). Respiratory tissues (nasal turbinate and lung) and brain were collected at 3 and 6 days after infection, and virus titers were quantified by performing plaque assays in Vero E6 cells stably expressing human TMPRSS2 (Vero E6/hTMPRSS2).

The transgenic hDPP4 hamsters supported MERS-CoV replication with infectious virus detected in the collected tissues (Fig. 3a). On day 3 after infection, the highest virus titer was in the nasal turbinate tissue (mean titer of 10^7.3^ pfu/g), while virus titers reached 10^4.9^, 10^3.7^, and 10^4.3^ pfu/g in the trachea, lung and brain tissues, respectively. By day 6 after infection, virus titers were reduced by 1000-fold in the trachea and nasal turbinate tissues, whereas titers increased by 10-fold in the lung tissue and 1000-fold in the brain.

**Figure 3.**
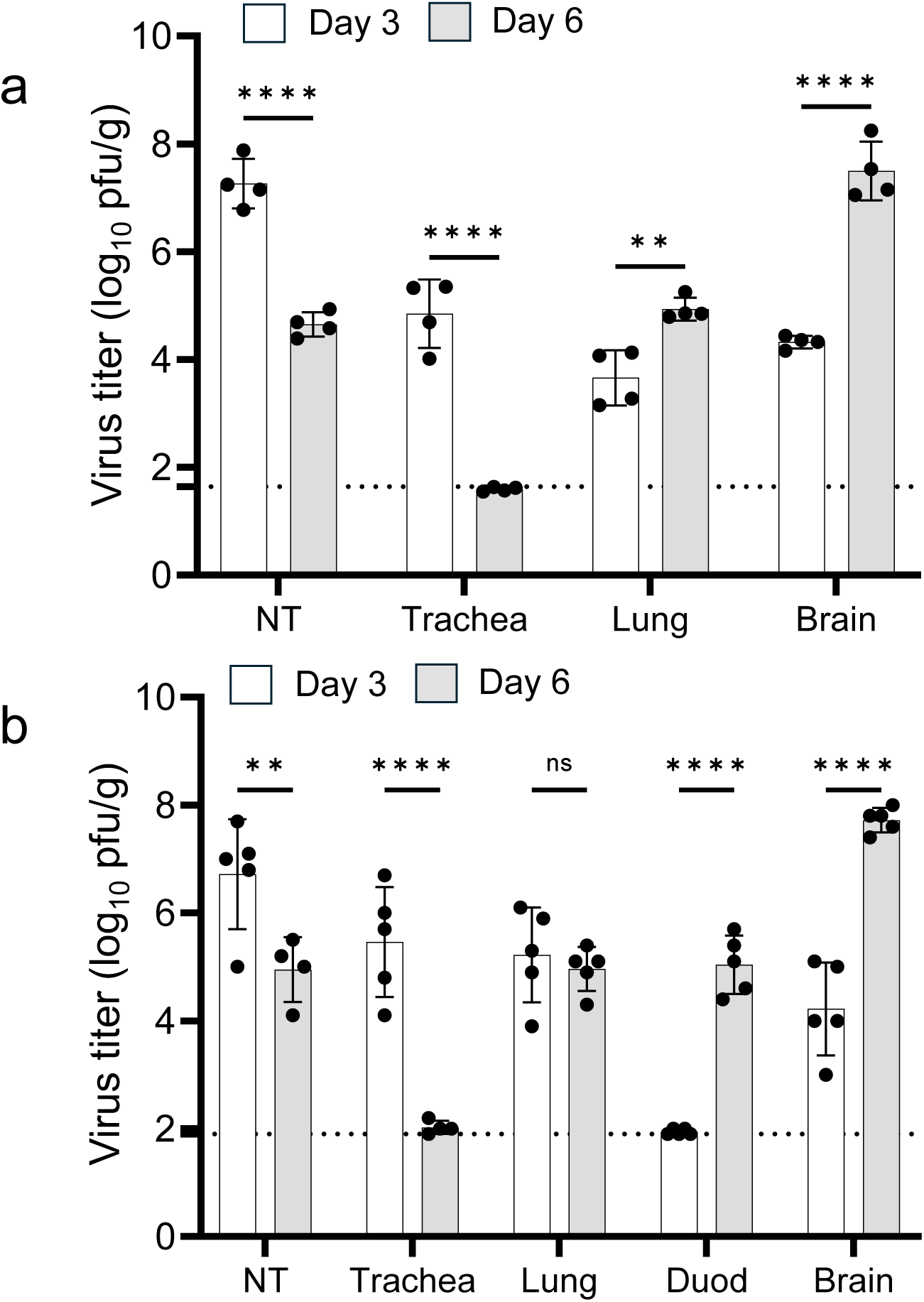
Transgenic hDPP4 hamsters support MERS-CoV replication. **a** Transgenic hamsters (n=4, males, 4–5 months old) were intranasally inoculated with 10^5^ pfu of MERS-CoV in a total volume of 30 µl of inoculum while under anesthesia. **b** A separate group of hDPP4 hamsters (n=5, females, 5–6 months old) were intratracheally inoculated with 10^5^ pfu of MERS-CoV in a total volume of 100 µl of inoculum while under anesthesia. In both groups, the indicated tissues were collected three days after infection and viral titers were determined by performing plaque assays with Vero E6/hTMPRSS2 cells. Data are means ± standard deviation with individual dots indicating data from individual hamsters. Data were analyzed by using a two-way analysis of variance (ANOVA). Vertical bars show the mean virus titer ± standard deviation with the lower limit of detection indicated by the horizontal dashed line. NT, nasal turbinate; Duod, duodenum; ns, not significant; pfu/g, plaque-forming units/gram. Data are from one experiment and source data are provided as a Supplementary Data file.

Next, hDPP4 hamsters (n=5, females, 5–6 months old) were inoculated with the same dose of virus, but via the intratracheal route (100 µl total volume). Similar to intranasal inoculation, the highest virus titer on day 3 after infection was in the nasal turbinate tissues (mean titer of 10^6.7^ pfu/g; Fig. 3b). In the trachea, virus replicated efficiently early after infection and was nearly cleared by day 6. In contrast, no detectable infectious virus was detected in gastrointestinal tissue (duodenum) on day 3 after infection but was detected at day 6 (mean titer of 10^5.0^ pfu/g). In the lung tissue, there was no significant difference in virus titers between the two timepoints; however, virus titers in the brain were 10,000-fold higher on day 6 after infection compared to the early time point.

Overall, these data demonstrate that hDPP4 transgenic Syrian hamsters support robust MERS-CoV replication in the respiratory tract with dissemination of the virus to other tissues.

### Evaluation of antiviral efficacy in hDPP4 transgenic hamsters against MERS-CoV

Having established that transgenic hDPP4 hamsters are susceptible to MERS-CoV infection and support efficient viral replication, we next evaluated their suitability as a small-animal model for antiviral drug testing. First, hDPP4 hamsters (n=8, females, 4–6 months old) were intranasally inoculated with 10^5^ pfu of MERS-CoV EMC/2012. Following a similar protocol used for wild type Syrian hamsters infected with SARS-CoV-2 (BA.2) ^18^, animals were then treated orally twice daily (at 12-h intervals) for three consecutive days with nirmatrelvir (200 mg/kg per dose, 300 µl inoculum; n=4) or vehicle control only (300 µl inoculum; n=4), beginning 24 h after infection. Tissues were collected four days after infection (12 h after the final treatment) for viral titration.

Treatment of MERS-CoV-infected hDPP4 hamsters with nirmatrelvir significantly reduced infectious virus titer by 3,162-fold in the lung tissue compared with vehicle control-treated animals (Fig. 4; *p*-value < 0.0001). In addition, virus titers in the trachea were significantly reduced (50-fold reduction; *p*-value = 0.006), and those in the nasal turbinate tissue were reduced, but not significantly (10-fold reduction; *p*-value = 0.087). However, no reduction in viral titer was observed in the gastrointestinal tissue (duodenum) and brain following drug treatment, suggesting limited drug penetration of nirmatrelvir in these tissues of hDPP4 hamsters.

**Figure 4.**
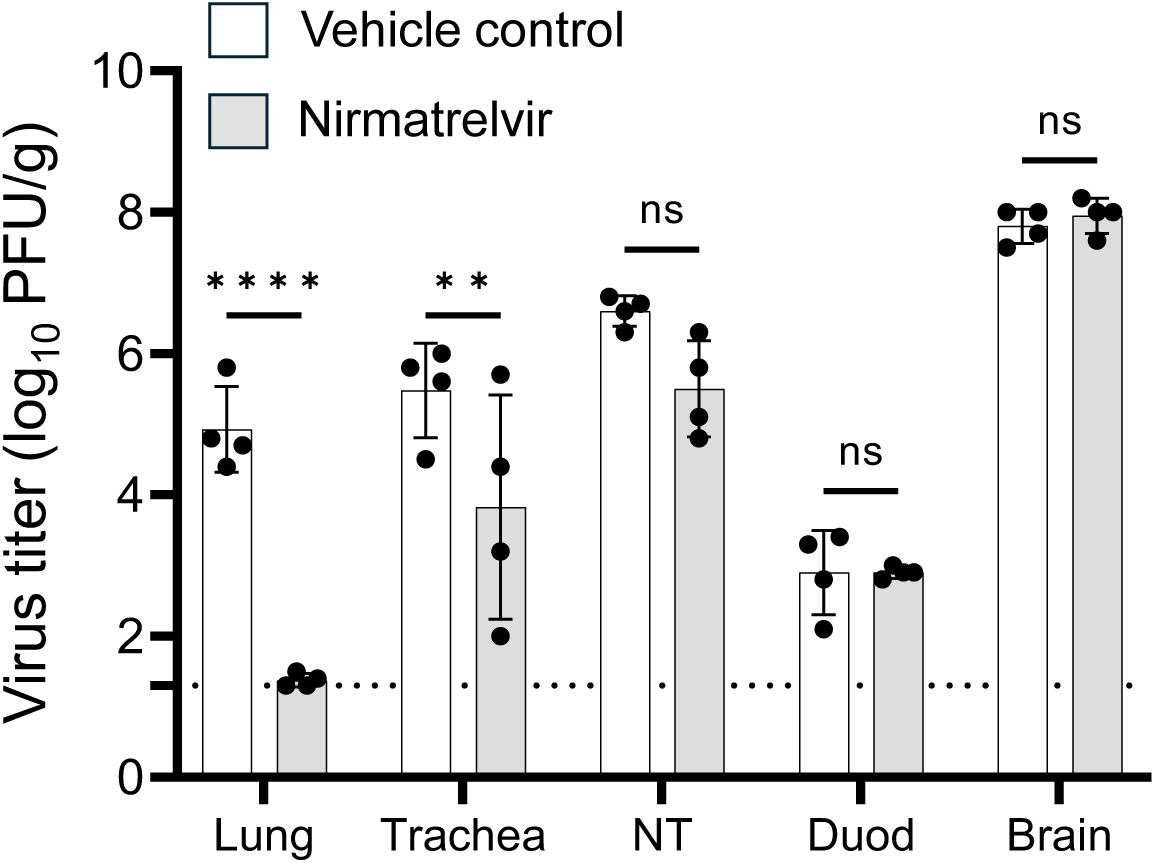
Therapeutic efficacy of the protease inhibitor nirmatrelvir against MERS-CoV in hDPP4 hamsters. Transgenic hamsters (n=8, females, 4–6 months old) were intranasally inoculated with MERS-CoV (10^5^ pfu in 30 µL total volume) while under anesthesia. Twenty-four hours after infection, animals were treated orally with either vehicle only (n=4, DMSO/PEG300; 300 µL total volume/dose) or nirmatrelvir (n=4, 200 mg/kg/dose; 300 µL total volume/dose) over three days with dosing every 12 h. The indicated tissues were collected four days after infection (12 h after the last treatment), and viral titers were determined by performing plaque assays with Vero E6/hTMPRSS2 cells. Data are means ± standard deviation with individual dots indicating data from individual hamsters. Data were analyzed by using a two-way analysis of variance (ANOVA). Vertical bars show the mean virus titer ± standard deviation with the lower limit of detection indicated by the horizontal dashed line. NT, nasal turbinate; Duod, duodenum; pfu/g, plaque-forming units/gram ns, not significant; **** *p*-value <0.0001; ** *p*-value = 0.006. Data are from one experiment and source data are provided as a Supplementary Data file.

### MERS-CoV transmits in hDPP4 hamsters by direct contact but not by airborne droplets

To evaluate the potential for airborne transmission in the absence of physical contact, we performed a respiratory droplet transmission experiment with hDPP4 hamsters as described previously ^19^. Donor hamsters (n=4, females, 4–6 months old) were intranasally inoculated with 10⁵ pfu of MERS-CoV EMC/2012, and one day after infection, naïve hamsters were placed in adjacent cages separated by a 5-cm wire mesh barrier to prevent direct contact while allowing airflow. Four days after infection or exposure, tissues from infected or contact hamsters and were analyzed for viral replication. While infectious virus was detected in the infected donor animals, no infectious virus was recovered from the contact animals after 72 h of exposure (Supplementary Fig. 1).

Next, we examined whether MERS-CoV transmitted to naïve hamsters by direct contact. Twenty-four hours after infection with 10^5^ pfu of virus, infected hamsters (n=5, females, 5–6 months old), paired in a clean cage with a naïve hamster (previous cage mate of the infected hamster). On day four after infection, the donor hamsters began to exhibit weight loss and were euthanized when they reached predefined criteria (Fig. 5; red lines). Seven days after exposure, the naïve contact hamsters also began to exhibit weight loss and were all euthanized by day 13 after exposure (day 14 after infection of the donor hamsters; Fig. 5; black lines). These findings establish that while airborne transmission of MERS-CoV was not detected under our experimental conditions, the virus is readily transmitted by direct contact in the hDPP4 hamster model similar to clinical observations of close-contact transmission in human cases ^20,21^.

**Figure 5.**
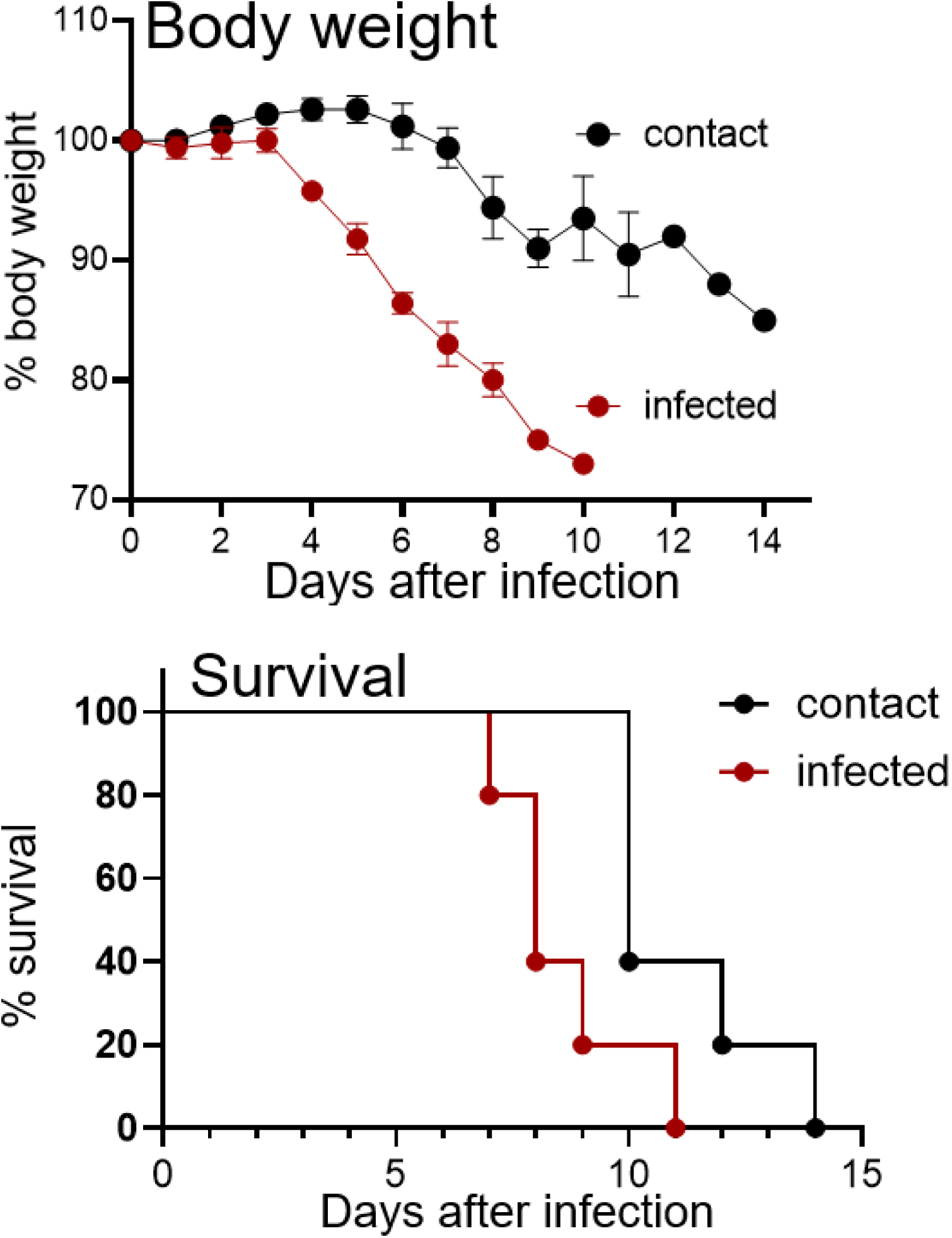
Direct contact transmission of MERS-CoV in hDPP4 hamsters. **a** Body weight and **b** survival were monitored for 14 days. Transgenic hamsters (n=5, females, 5–6 months old) were infected with MERS-CoV (10^5^ pfu in 30 µL total volume) while under anesthesia. Twenty-four hours after infection, infected animals were moved to a cage with naïve, contact hDPP4 hamsters and **a** body weight and **b** survival were monitored for 14 days. Data are mean percentages of starting weights ± standard deviation. Survival data were analyzed by using the log-rank (Mantel–Cox) test.

### Vaccination of hDPP4 hamsters reduces disease severity following direct contact transmission

Finally, we assessed the efficacy of vaccination of hDPP4 hamsters to reduce virus replication and transmission. One group of hamsters (n=4, males, 6–7 months old) was immunized intramuscularly twice (4-weeks apart) with 10 µg of purified MERS receptor-binding domain (RBD) antigen with alum while a control group (n=4) was mock-vaccinated (alum only). Four weeks after the second immunization, both groups of hamsters were infected with 10^5^ pfu of MERS-CoV EMC/2012. At three days after infection, virus titers in the lung tissue of RBD-vaccinated hamsters were significantly reduced (X-fold; *p*-value = 0.0009) compared to control animals (Fig. 6a). Virus titers were also reduced in other tissues (trachea, nasal turbinate, and brain), but not significantly.

**Figure 6.**
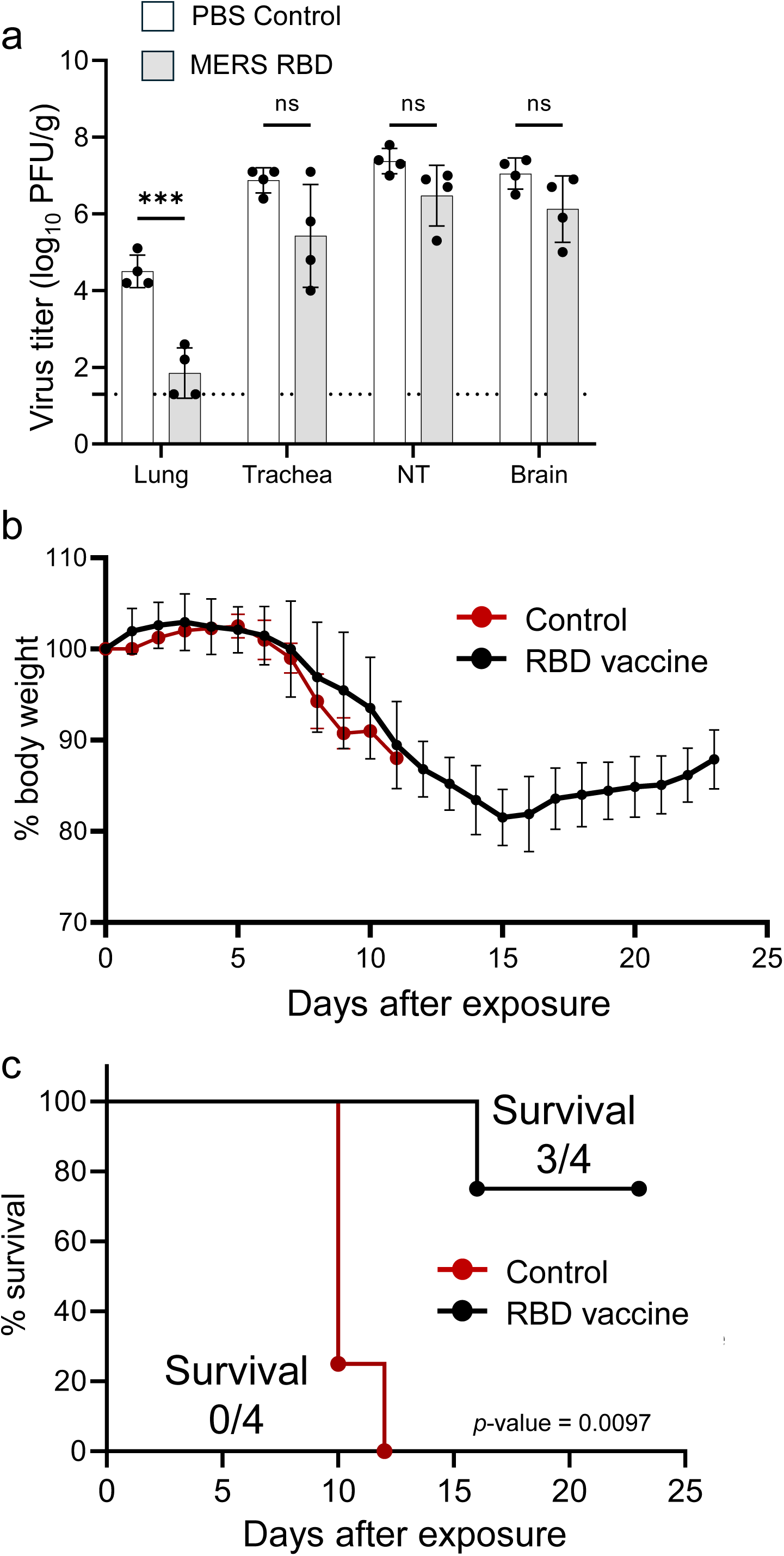
Immunization with the MERS-RBD reduces virus replication in the lung tissue and disease severity but not direct contact transmission efficiency. Transgenic hamsters (males, 6–8 months old) were immunized twice (28-days apart) intramuscularly with 10 µg of purified MERS-RBD plus alum or with alum only. **a** To determine virus replication after immunization, hDPP4 hamsters were intranasally inoculated 28 days after the final immunization with MERS-CoV at a dose of 10^5^ pfu (30 µl total volume of inoculum; n=4 RBD+alum immunized and n=4 alum only) while under anesthesia. Virus replication was examined three days after infection in the indicated tissues by performing plaque assays with Vero E6/hTMPRSS2 cells. Data are means ± standard deviation with individual dots indicating data from individual hamsters and were analyzed by using a two-way analysis of variance (ANOVA). Vertical bars show the mean virus titer ± standard deviation with the lower limit of detection indicated by the horizontal dashed line. NT, nasal turbinate; pfu/g, plaque-forming units/gram ns, not significant; *** *p*-value = 0.0009. To examine direct contact transmission, non-immunized, infected hDPP4 hamsters (n=8) were paired with contact animals that were either immunized with purified MERS RBD plus alum (n=4) or alum only (n=4). After pairing, hDPP4 contact hamsters were monitored for 24 days for **b** weight loss and **c** survival. Data are mean percentages of the starting weights ± standard deviation. Survival data were analyzed by using the log-rank (Mantel–Cox) test. Data are from one experiment and source data are provided as a Supplementary Data file.

To determine whether immunization with purified MERS RBD could prevent direct contact transmission, hDPP4 hamsters (n=8, males, 7–8 months old) were infected with 10^5^ pfu of MERS-CoV EMC/2012. Twenty-four hours after infection, the infected donor hamsters were moved to the cage of a contact hDPP4 hamster (previous cage mates). Infected hDPP4 hamsters were paired with either hDPP4 contact hamsters that were intramuscularly immunized twice with the MERS RBD antigen (10 µg of antigen with alum; n=4) or mock-immunized (alum only; n=4). Both groups of contact hamsters exhibited body weight loss on days 5–6 after infection (Fig. 6b) with the mock-immunized hamsters succumbing to infection by day 11 after exposure (Fig. 6c). In contrast, only one immunized hDPP4 contact hamster succumbed to infection (on day 16 after exposure) (Fig. 6c), demonstrating that vaccination with the MERS RBD antigen reduces disease severity (75% survival; *p*-value = 0.0097) but does not reduce direct contact transmission as evident by weight loss in all MERS RBD-immunized contact hamsters. In the one RBD-immunized contact animal that did not survive, infectious virus was detected in respiratory (nasal turbinate, trachea, and lung) and brain tissues (Supplementary Fig. 2).

Taken together, our findings demonstrate that hDPP4 transgenic Syrian hamsters are a versatile platform for the preclinical evaluation of antiviral therapeutics and for mechanistic studies of MERS-CoV pathogenesis and transmission.

## Discussion

Syrian hamsters were a valuable animal model during the COVID-19 pandemic to examine and compare the pathogenicity and transmission of different SARS-CoV-2 variants and evaluate medical countermeasures against the emerging virus^22,23^. However for MERS-CoV, wild type Syrian hamsters do not support viral infection due to species-specific receptor incompatibility^24,25^. Therefore, nonhuman primates such as rhesus macaques and marmosets have been used to model MERS-CoV infection. In rhesus macaques, infected animals develop mild disease with increased in body temperature but no weight loss. Viral replication has been detected in the lower but not upper respiratory tract^26^. In contrast, infection of marmosets with MERS-CoV results in a more severe disease, with some animal succumbing to lethal pneumonia^27^. MERS-CoV replicates efficiently to high virus titers in the respiratory tract tissues of marmosets. Interestingly, viral titers in the lung tissues of marmosets do not decrease between days 3 and 6 after infection, an observation similar to what we observed in our studies with transgenic hDPP4 hamsters. Virus dissemination is prominent in MERS-CoV–infected marmosets, with virus or viral RNA detectable in multiple tissues, including the liver, spleen, brain, and gastrointestinal tract. Although the breadth of tissue sampling was limited in our study, we likewise detected infectious virus at extrapulmonary sites, including the duodenum and brain.

For small animal modeling of MERS-CoV, multiple transgenic mouse models expressing hDPP4 have been developed. These include lines in which hDPP4 is expressed under the endogenous mouse promoter, as well as models driven by heterologous human promoters (e.g., keratin 18, the same promoter we used in hDPP4 hamsters), and have been widely used for pathogenesis and countermeasure studies^28-30^. In the transgenic hDPP4 mouse models, infected animals experience weight loss and succumb to infection typically between days 6 to 7 after infection, a similar disease outcome to that seen in hDPP4 hamsters. Virus replication in the lung tissue of transgenic mice is robust, with virus titers reaching 10^7^–10^8^ infectious virus particles in tissue depending on the assay^29,30^.

Direct contact and exposure of infected dromedary camels can lead to human cases of MERS-CoV with resulting onward human-to-human transmission, although continuous transmission in humans is not sustained, unlike SARS-CoV-2^31^. Despite efficient viral replication in the lungs of transgenic mice, MERS-CoV transmission in this small-animal model is limited^30^, representing a major drawback of this animal model. In contrast, our hDPP4 hamster model supports robust direct contact transmission of MERS-CoV, providing an improved platform to examine transmission dynamics and evaluate medical countermeasures.

We successfully used this transgenic hamster model to evaluate the efficacy of a standard vaccine antigen (purified MERS-CoV RBD) against infection. Although RBD-immunized contact hDPP4 hamsters were not fully protected against infection following direct transmission from infected donor hamsters, most of the vaccinated animals survived. Because immunization was delivered intramuscularly, limited induction of respiratory mucosal immunity may have contributed to the incomplete protection against infection^32,33^, an important limitation that likely constrained the full exploitation of this model’s utility for transmission studies. Future work should explore intranasal vaccination strategies to better elicit mucosal immunity and improve protection in the upper respiratory tract.

The primary objective of this work was to establish a transgenic hDPP4 hamster model with robust viral replication and transmission, and to demonstrate the utility of this model for evaluating medical countermeasures. A key limitation is the lack of detailed pathological analyses. Future studies will characterize the pulmonary disease phenotype and identify the specific lung cell types targeted by infection, enabling direct comparison with the pathology and tropism observed in human MERS-CoV infection.

Collectively, our findings demonstrate that transgenic hDPP4 transgenic Syrian hamsters are a new, useful tool for MERS-CoV research. As a small-animal model, they support the efficient infection of both the upper and lower airways, allow close-contact transmission, and permit medical countermeasure testing. This model will enhance our ability to investigate coronavirus pathogenesis and develop countermeasures against current and emerging zoonotic coronaviruses that use hDPP4 as a cellular receptor.

## Methods

### Animal experiments and approvals

Transgenic hDPP4 hamsters were generated at Utah State University (USU) under protocol #13758, which was approved by the USU Institutional Animal Care and Use Committee. Infection studies of hDPP4 hamsters were performed at the University of Wisconsin-Madison (UW) in accordance with the UW Institutional Animal Care and Use Committee under protocol V006426.

Animals were housed in a temperature- and humidity-controlled environment with 14-h light/10-h dark cycles for breeding (USU) and 12-h light and dark cycles (UW) for infection studies. Prior to the start of a study, hamsters were allowed to acclimate to the animal room for at least three days. Enrichment was provided and water and food were provided *ad libitum*.

Sample sizes were determined based on previous animal studies with other coronaviruses; no power calculations were performed prior to the studies. Animals were randomly taken off the ventilated caging rack and placed into the study. Because the studies were performed under BSL 3 AG containment and samples were inventoried, samples were not blinded to the researchers.

### Construction of the Gateway expression vector pmhyGENIE-3-K18-hDPP4

The entry clone containing the K18-hDPP4-T2A-GFP cassette was generated through a series of DNA subcloning steps. First, vector pKO2.1-hDPP4-T2A-GFP was constructed by NEBuilder HiFi DNA assembly (NEB, E2621). Briefly, human DPP4 CDS, the T2A self-cleaving linker, and GFP were PCR-amplified with 25-bp overlaps at each end and then assembled with linearized plasmid pKO2.1. To introduce the K18 promoter and other regulatory sequences, plasmid pKB12 (containing the K18 5’ and 3’ regions) and pKO2.1-hDPP4-T2A-GFP were digested with XbaI and NotI, followed by ligation to produce pKB12-K18-hDPP4-T2A-GFP. Next, pKB12-K18-hDPP4-T2A-GFP was digested with HpaI and BamHI and the resulting K18-hDPP4-T2A-GFP fragment was ligated to pENTR^TM^-1A to generate the Gateway entry vector pENTR^TM^-1A-K18-hDPP4-T2A-GFP.

The Gateway destination vector pmhyGENIE-3 (piggyBac-based transposon system), was generously provided by Stefan Moisyadi (University of Hawaii). The expression plasmid used in pronuclear (PN) injection, pmhyGENIE-3-K18-hDPP4, was constructed by Gateway cloning system (Invitrogen, Cat.11791019) per the manufacturer’s instructions. The donor vector, pENTR^TM^-1A-K18-hDPP4-T2A-GFP, was recombined with pmhyGENIE-3 by LR Clonase enzyme to generate the expression vector pmhyGENIE-3-K18-hDPP4. Plasmid pmhyGENIE-3-K18-hDPP4 was confirmed by Sanger sequencing before use in pronuclear injection.

### Generation of transgenic hDPP4 golden Syrian hamster by PN injection

Golden Syrian hamsters were obtained from Charles River and bred in house. Embryo manipulation and PN injection were performed as described previously^34^ ^35^. Briefly, female golden Syrian hamsters were superovulated by i.p. injection of 10–25 IU of PMSG (BioVendor, Cat.RP1782725000) based on body weight in the morning (9–12 AM) on Day 1 of the estrous cycle. These females were mated with fertile males after 6:00 PM on Day 4 of the estrous cycle and were euthanized approximately 18 h after mating for zygotes isolation. Zygotes were flushed from oviducts with warmed, equilibrated HECM-9 medium supplemented with 0.5 mg/mL human serum albumin. Embryos were then washed twice with HECM-9, transferred into 20-µL of HECM-9, covered by mineral oil, and cultured at 37.5°C under 10% CO_2_, 5% O_2_, and 85% N_2_.

The plasmid pmhyGENIE-3-K18-hDPP4 was diluted to 8 ng/uL using pH7.0 TE buffer. All procedures were performed in a dark room, and red filters were used for all microscope light sources. To perform PN injections, groups of 15–20 hamster zygotes were transferred to a 100-µL HECM-9 drop in the microinjection dish and the plasmid was injected into the male pronucleus indicated as PN swollen. After injection, embryos were washed twice with equilibrated HECM-9 and cultured at 37.5°C under 10% CO_2_, 5% O_2_, and 85% N_2_ until embryo transfer. Embryos with normal morphology were selected for transfer into the oviducts of pseudopregnant recipients that were prepared by matting with vasectomize males one day previously (at the same time as the zygote donors were mated). Embryos were transferred bilaterally with 10–15 embryos per oviduct. To genotype founder and subsequent generation transgenic hamsters, genomic DNA (gDNA) was isolated from toe clippings using the Qiagen Blood and Tissue Kit (Qiagen, Cat.69506). PCR for amplifying overlap hDPP4 F11-R11 and overlap hDPP4 F31-R31 was performed with LA *Taq* (Takara, Cat. RR002A) and the following parameters: initial denaturation at 94°C for 1 min followed by 32 cycles of 10 s denaturation at 98°C, and 4 min extension at 68°C, with a final extension step of 72°C for 10 min. PCR for amplifying overlap hDPP4 F26-R26 was performed with Q5 with high GC buffer (NEB, Cat.M4093) and the following parameters: initial denaturation at 98°C for 30 s followed by 32 cycles of 10 s denaturation at 98°C, 30 s annealing at 68°C, and 90 s extension at 72°C, with a final extension step of 72°C for 5 min. Primers used for genotyping transgenic pups are summarized in Table 1.

### Characterization of the hDPP4 transgene integration site by TLA

The hDPP4 transgene integration site was characterized using Targeted Locus Amplification (TLA) by Cergentis (Utrecht, The Netherlands). Junction-specific primers based on the 5’ and 3’ breakdown sequences are listed in Table 2. PCR for amplifying the genomic-transgene junction was performed with Ex Taq (Takara, Cat.RR001A) under the following parameters: initial denaturation at 94°C for 30 s followed by 32 cycles of 10 s denaturation at 98°C, 30 s annealing at 60°C, and 30 s extension at 72°C, with a final extension step of 72°C for 5 min.

### Biosafety containment

All infection experiments with MERS-CoV were performed under biosafety level 3 agriculture (BSL3 AG) containment at the University of Wisconsin-Madison, which is approved for such use by the Centers for Disease Control and Prevention and by the US Department of Agriculture.

### Cells and virus

Vero E6/hTMPRSS2 cells (JCRB 1819) were cultured in Dulbecco’s modified Eagle’s medium (DMEM) containing 10% fetal bovine serum (FBS), 100 U/ml penicillin-streptomycin, and 1 mg/ml geneticin (G418; Invivogen). MERS-CoV strain EMC/2012 was propagated on Vero E6/hTMPRSS2 cells. Virus stocks were aliquoted and stored at −80°C until use.

### Experimental infection of hDPP4 hamsters and sample collection

Transgenic hDPP4 hamsters (sex and age are indicated in the Results section) were infected either by intranasal or intratracheal inoculation with 10^5^ pfu of virus in a total volume of 30 µl or 100 µl, respectively, while under isoflurane anesthesia. For tissue collection, animals were humanely euthanized by an isoflurane overdose and cervical dislocation a second method to verify death. Tissues were collected in duplicate and frozen at -80°C for at least 24 h. Virus titers were determined by performing plaque assays on Vero E6/hTMPRSS2 cells overlaid with 1% methylcellulose solution.

### Antiviral treatment and vaccination

Nirmatrelvir (PF-07321332; MedChemExpress) was suspended in DMSO (10%) and then added to PEG300 (MedChemExpress). Hamsters were treated with 200 mg/kg nirmatrelvir twice daily (12-h intervals) given orally via an 18-gauge feeding tube. Control animals were treated with vehicle only. Hamsters were treated 24 h after infection for three consecutive days. Twelve hours after the last treatment, animals were humanely euthanized and tissues were collected.

Hamsters were immunized twice (28-weeks apart) with 10 µg of purified MERS receptor-binding domain protein (R&D Systems) along with 2% Alhydrogel (InvivoGen; final concentration of 0.45 µg of alum). Animals were immunized by intramuscular inoculation in the leg with a total volume of 50 µl. Control animals were inoculated with only Alhydrogel. After the final immunization, hDPP4 hamsters were directly infected with MERS-CoV or served as contact animals in a direct transmission study.

### Transmission experiments

For transmission studies, hamster pairs were previously cage mates to reduce stress and prevent fighting. Hamsters were separated for 24 h after infection and then paired with the original cage mate for the transmission study. For direct contact transmission studies, the infected hDPP4 hamsters were always moved to the cage of the contact animals.

For the airborne transmission study, donor hamsters were housed in the same cage as the infected hamsters but were separated by a 5-cm wire-mesh barrier that allowed airflow but prevented physical contact. Cages were placed in the transmission unit with directional airflow from the infected to the contact animal.

### Statistical analysis

GraphPad Prism software was used to analyze the virus titer data from the lung and nasal turbinate tissues as well as the weights of the hamsters. Kaplan–Meier survival curves were analyzed by using the log-rank (Mantel–Cox) test. Data are presented as means ± standard deviation (SD).

## Data availability

The source data behind the graphs in the paper can be found in Supplementary Data 1.

## Acknowledgments

This work was supported by the National Institute of Allergy and Infectious Diseases of the National Institutes of Health under Award Number P01AI165077 to PJH. We thank Sue Watson for editing the manuscript.

## Author contributions

Conceptualization: ZW and PJH

Methodology and data analysis: TW, YL, RL, NM, ZW, and PJH

Investigation: TW, YL, RL, NM, and PJH

Visualization: TW, YL, RL, ZW, and PJH

Writing – original draft: TW, YL, ZW, and PJH

Writing – review & editing: TW, YL, RL, NM, ZW, and PJH

## Supplementary Figures

**Supplementary Figure 1. Virus titers from infected, donor hDPP4 hamsters.** Donor hamsters (n=4, females, 4–6 months old) were intranasally inoculated with 10⁵ pfu of MERS-CoV EMC/2012 and three days after infection, the indicated tissues were collected to examine virus titers. Data are means ± standard deviation with individual dots indicating data from individual hamsters. Vertical bars show the mean virus titer ± standard deviation, with the lower limit of detection indicated by the horizontal dashed line. NT, nasal turbinate; pfu/g, plaque-forming units/gram. Data are from one experiment and source data are provided as a Supplementary Data file.

**Supplementary Figure 2. Virus titers from the MERS-RBD immunized hDPP4 hamster that succumbed to infection.** Virus titers from the indicated tissues of the immunized hamster that succumbed to infection on day 16 after exposure to a infected, donor hamster. The lower limit of detection is indicated by the horizontal dashed line. NT, nasal turbinate; pfu/g, plaque-forming units/gram. Data are from one experiment and source data are provided as a Supplementary Data file.

