## Supplementary figures and images for "Transgenic human dipeptidyl peptidase-4 Syrian hamsters support MERS coronavirus infection and contact transmission"

### Supplemental Figures 1-2

#### Supplementary Figure 1

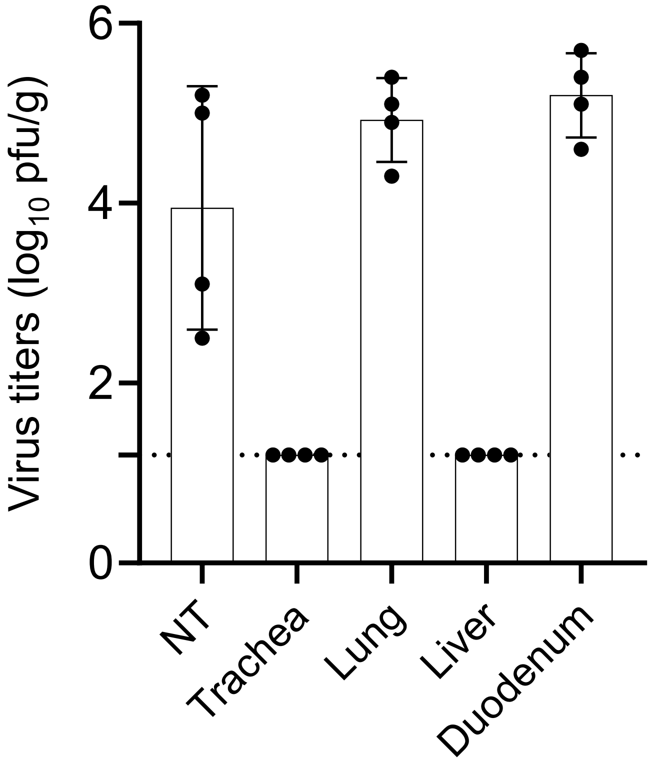

#### Supplementary Figure 2

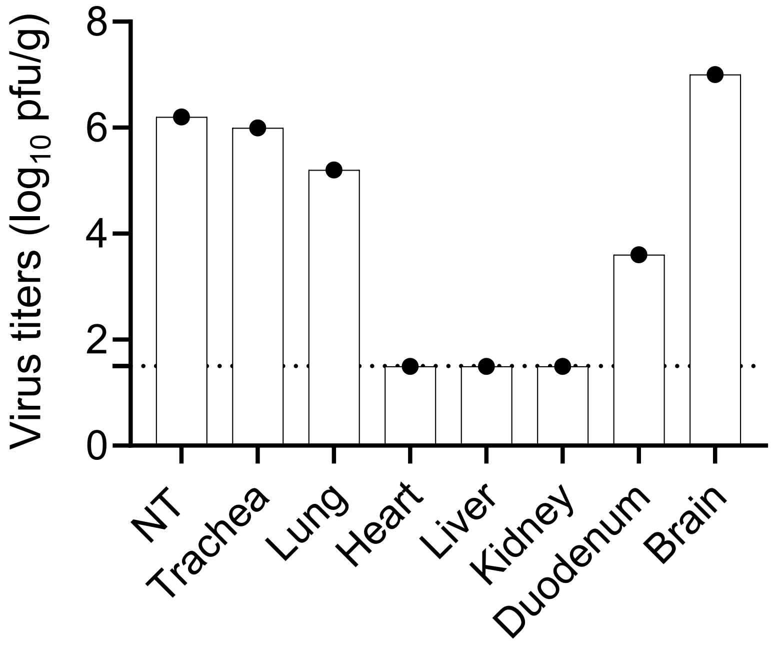
